# Network Analysis of Inflammatory Bowel Disease Reveals PTPN2 As New Monogenic Cause of Intestinal Inflammation

**DOI:** 10.1101/768028

**Authors:** Marianna Parlato, Julia Pazmandi, Qing Nian, Fabienne Charbit-Henrion, Bernadette Bègue, Emmanuel Martin, Marini Thian, Felix Müller, Marco Maggioni, Rémi Duclaux-Loras, Frederic Rieux-Laucat, Thierry-Jo Molina, Sylvain Latour, Frank Ruemmele, Jörg Menche, Fernando Rodrigues-Lima, Kaan Boztug, Nadine Cerf-Bensussan

**Author notes:** to whom correspondence should be addressed: Nadine Cerf-Bensussan or Kaan Boztug. Contributed equally (Marianna Parlato, Julia Pazmandi, Qing Nian, Fernando Rodrigues-Lima, Kaan Boztug, Nadine Cerf-Bensussan). **Author contributions:** JP, MP, JM, KB, NC-B. study design and manuscript writing. MP, JP, FM, NQ, BB, EM, MM, RDL, FR-L TJM: data acquisition. FC-H, FR: patient care coordination and clinical samples acquisition. MP, JP, FL-R, SL, FM and MT analysis and interpretation of data. FL-R, KB, NC-B: Study supervision. The manuscript was reviewed and approved by all co-authors.

## Abstract

**BACKGROUND & AIMS:** Genome-wide association studies (GWAS) have uncovered multiple loci associated with inflammatory bowel disease (IBD), yet delineating functional consequences is complex. We used a network-based approach to uncover traits common to monogenic and polygenic forms of IBD in order to reconstruct disease relevant pathways and prioritize causal genes.

**METHODS:** We have used an iterative random walk with restart to explore network neighborhood around the core monogenic IBD cluster and disease-module cohesion to identify functionally relevant GWAS genes. Whole exome sequencing was used to screen a cohort of monogenic IBD for germline mutations in top GWAS genes. One mutation was identified and validated by a combination of biochemical approaches.

**RESULTS:** Monogenic IBD genes clustered siginificantly on the molecular networks and had central roles in network topology. Iterative random walk from these genes allowed to rank the GWAS genes, among which 14 had high disease-module cohesion and were selected as putative causal genes. As a proof of concept, a germline loss of function mutation was identified in *PTPN2,* one of the top candidates, as a novel genetic etiology of early-onset intestinal autoimmunity. The mutation abolished the catalytic activity of the enzyme, resulting in haploinsufficiency and hyper-activation of the JAK/STAT pathway in lymphocytes.

**CONCLUSIONS:** Our network-based approach bridges the gap between large-scale network medicine prediction and single-gene defects and underscores the crucial need of fine tuning the JAK/STAT pathway to preserve intestinal immune homeostasis. Our data provide genetic-based rationale for using drugs targeting the JAK/STAT pathway in IBD.

## INTRODUCTION

One of the prototypic diseases at the crossroads of autoimmunity and autoinflammation is inflammatory bowel disease (IBD)^1^. IBD is commonly divided into ulcerative colitis and Crohn’s disease, denoting a group of heterogenous, chronic disorders of the gastrointestinal tract that affect approximately 1/1,000 individuals in Western countries with a peak onset in young adults^2^. It is now acknowledged that IBDs result from pathologic interactions between the microbiota and the host immune system in genetically susceptible individuals^3^. The contribution of genetics to these diseases of relatively late onset is however weak with more than 230 identified susceptibility loci, which, altogether, account for only 15% of IBD risk^4–6^. To date, common variants associated with IBD cannot predict IBD susceptibility and disease progression with high confidence for individual patients^7^. Accordingly, much of the heritability of IBD remains unexplained^8^ and one of the major challenges of today is linking significantly associated genes through GWAS studies to the pathobiology of IBD. Interestingly, IBD can also manifest as an early-onset monogenic disease^9^. While to date only few gene defects have been found to cause monogenic IBD affecting exclusively or predominantly bowel immune homeostasis, IBD-like features can be the first signs of an underlying, systemic inborn errors of immunity as illustrated in over 50 Mendelian disorders that can present with an IBD-like phenotype^9^. These monogenic forms of IBD point out to crucial non redundant molecular mechanisms maintaining intestinal homeostasis, both involving innate and adaptive immunity, while GWAS typically yield many genes with moderate effect size and often unclear pathobiological impact. In light of recent discoveries identifying diseases as localized clusters on networks^10,11^, we here set out to leverage the network-based characteristic of monogenic IBD to gain more understanding of the biological context of IBD-associated GWAS data, and to bridge the gap between the adaptive (monogenic) and innate immune component of IBD. We have used a state-of-the art network-based approach to uncover novel traits of both monogenic and common polygenic forms of IBD with the aims of reconstructing disease-relevant pathways and prioritizing causal genes. Our network-based approach predicted PTPN2 as an important regulator of intestinal immune homeostasis, a hypothesis also supported by data derived from mouse studies ^12–14,15^. Further demonstrating the central role of PTPN2, we identified *PTPN2* haploinsufficiency as a novel, monogenic inborn error of immunity impairing intestinal immunoregulation, bridging the gap between large-scale network medicine-based prediction and single-gene defects and the dissection of the underlying molecular pathomechanisms.

## METHODS

### Gene Sets

Monogenic defects underlying IBD were defined as recently described^9^. We have opted to use a widely-utilized and cited resource to obtain our list of GWAS genes, therefore the GWAScatalog^16^ was used to query significant SNPs associated with IBD by queries for “inflammatory bowel disease”, “Crohn’s disease” and “ulcerative colitis” respectively (https://www.ebi.ac.uk/gwas/).

### Molecular Interaction Networks And Network-Based Measures

A full description of all network-based is provided in the Supplementary Materials and Methods. In brief, we collected a total of 22 molecular networks including <u>a</u> manually curated interactome network of physical, experimentally validated protein-protein interactions which consists of 13,460 proteins and 141,296 interactions^10^, and 16 transcriptome-wide coexpression networks from^17^.We constructed pathway similarity networks using the Molecular Signatures Database (MSigDB, http://software.broadinstitute.org/gsea/msigdb/collection_details.jsp). In each pathway similarity network, two genes were connected if they shared a pathway annotation. The number of shared pathways determined the weight of an edge.

We calculated network-based separation of two diseases A and B as defined by:

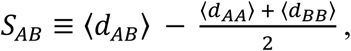

where *s*_AB_ compares the shortest distances between proteins *within* each disease, 〈*d_BB_*〉 and 〈*d_AA_*〉, to the shortest distances 〈*d_AB_*〉 *between* A-B protein pairs. Proteins associated with both A and B have *d*_AB_ = 0.

Gene prioritization was done using an iterative version of the random walk with restart^18^. The top candidates were selected based on disease-module cohesion, which is a property reflecting the localization of candidate genes around the initial monogenic gene sets detailed further in the Supplement.

### Gene Set Enrichments

To assess enrichment of proteins/genes of interest, the Fisher’s exact test was used to get odds ratio and p-value. The OrphaNet dataset for orphan genes from the Disgenet database (http://www.disgenet.org) was used to obtain a list of disease genes. The list of dispensable genes was extracted from the publication by Saleheen et al^19^. The list of essential genes was obtained from the online gene essentiality database (OGEE http://ogee.medgenius.info/browse/). The list of genes associated with OMIM traits was extracted from Menche et al^10^. Pathway enrichment was done using Enrichr ^20^ and visualized with a custom R script.

### Study Approval

All participants provided informed consent in accordance with the Declaration of Helsinki under institutional review board-approved protocol (CPP Ile-de-France II).

### DNA Sequencing

Whole-exome and targeted panel sequencing were performed as previously described^21^. For conventional Sanger sequencing, the PTPN2 gene was amplified by PCR from genomic DNA using the following primers: for-5-GGAGTCCCTGAATCACCAGC-3; rev-5-TTCACTGCCAGTGGAAGCAA-3.

### Variant Prioritization of WES Data

For WES analysis, candidate variants were ranked by filtering out common polymorphisms reported in public databases, including the Genome Aggregation Database (gnomAD)] http://gnomad.broadinstitute.org/ or seen in the 13,465 exomes sequenced at Institut Imagine from families affected with genetic diseases. Polyphen2 (http://genetics.bwh.harvard.edu/pph2/), Sift (http://sift.bii.a-star.edu.sg/) and Mutation Taster (http://www.mutationtaster.org/) were used to predict consequences of mutations on protein function. Mutations were next ranked on the basis of the predicted impact of each variant by combined annotation-dependent depletion (CADD), and compared with the mutation significance cutoff (MSC) (http://pec630.rockefeller.edu:8080/MSC/), a gene-level specific cutoff for CADD scores^22^.

### Western Blotting

Cell lysates were generated and Western blotting was performed as described in the Supplementary Materials and Methods.

### PTPN2 Immunoprecipitation and Activity Assay

HEK293T cells were transfected with FLAG-tagged PTPN2, PTPN2 C216G or empty vector (Origene). 24h after transfection, cells were washed with PBS and resuspended in lysis buffer (PBS, 0.5% Triton X-100, protease inhibitors cocktail). After centrifugation (15,000 g for 10 min at 4°C), supernatant was taken and total protein concentration measured with Bradford reagent. For PTPN2 immunoprecipitation, 1 mg of whole-cell extracts (200 μl) were incubated overnight with 1 μg of an anti-flag mouse monoclonal antibody (Origene) at 4 °C. Samples were then rocked for two hours at 4° C in presence of 20 μl protein G– agarose (Santa Cruz). The beads were harvested by centrifugation, washed 3 times with lysis buffer and incubated with 400 μL of phosphatase buffer containing 50 μM FAM-pStat1 peptide at 37 °C. After 30 min incubation, beads were spun down by centrifugation (2000 x g, 5 min) and 50 μl of supernatant were taken and mixed with 50 μL of HClO_4_ (15% in water) prior to RP-UFLC analysis as described previously ^23^. Beads were harvested and boiled in 40 μL of Laemmli sample buffer prior to western blot analysis with a polyclonal rabbit anti-PTPN2 antibody (SAB4200249, Sigma-Aldrich).

### Modeling of PTPN2 C216G Mutation

The PTPN2 C216G mutation was created and analyzed using Swiss-PdbViewer^24^ with human PTPN2 structure as template (PDB entry: 1L8K). Structures were rendered using Chimera^25^.

### Protein Sequence Alignment

Multiple sequences were aligned using ClustalW2^26^ (https://www.ebi.ac.uk/Tools/msa/clustalo/). Protein sequences were obtained from NCBI. Sequence alignment is based on the following RefSeq accession numbers: NP_536347.1, NP_446442.1, NP_001120649.1, NP_997819.1, NP_001030508.1, AFH27310.1, NP_001086910.1; NP_002818.1; NP_002826.3; NP_002820.3; NP_002829.3; NP_002824.1.

### Statistical Analysis

Results were analyzed with Prism (version 5.00; GraphPad software, Inc.) and p-values <0.05 were considered significant.

Additional methods are provided as Supplementary Material.

## RESULTS

### Monogenic IBDs form a connected cluster on biological networks

Genes associated with the same disease tend to form connected subgraphs, or *disease modules* within molecular interaction networks. This, in turn, can be used to design network-based algorithms for the identification of key implicated pathways and to predict novel disease-associated genes. To apply these approaches to monogenic IBD, a set of the 70 single-gene defects known to underlie bowel inflammation was established based on our recent review of the literature^9^ (Supplementary Figure 1A, Supplementary Table1). The majority of the respective gene defects cause disease through loss of function (Supplementary Figure 1B), often affecting adaptive immunity, most frequently T-cell function (20%, Supplementary Figure 1C). The strong enrichment in adaptive immune genes in monogenic IBD is in keeping with a recent analysis by Fischer and Raussel stressing the limited genetic redundancy in the adaptive immune system^27^. We constructed a comprehensive set of protein-protein interaction (ppi), co-expression and functional similarity networks, collectively reflecting the complex and interconnected physical, regulatory and functional systems within the cell and organism (Figure 1A). We found that monogenic IBD genes generally represented important nodes in the architecture of these molecular networks, illustrated by high connectivity, centrality and clustering coefficient scores on all network levels (Figure 1B-D, Supplementary Figure 2A-C). The connectedness of the 70 monogenic IBD genes on the different networks was further assessed by measuring the respective largest connected components (lcc) (Figure 1E). Monogenic IBD formed a significant disease cluster on a wide-range of biological networks (Figure 1F). The clustering was particularly pronounced in coexpression networks, indicating that genes responsible for inflammatory signals (many of these representing monogenic IBD genes) tend to be transcriptionally linked, as has been observed previously^4,28^. Overall, the connectivity of monogenic IBD was best captured by a manually curated interactome of physical protein-protein interactions, which consists of 13,460 nodes and over 141,296 connections (Figure 1E, Supplementary Figure 2B). To investigate the relationship between monogenic IBD and other complex traits, we calculated distance measures between monogenic IBD and other disease-associated genes on the interactome (Figure 1G). Monogenic IBD was in closest proximity (as measured by the separation parameter *s*_AB_) to both “primary immunodeficiencies” and “gastrointestinal diseases” (Figure 1G) as compared to other immune-mediated diseases such as rheuma and type I diabetes (Figure 1H). Overall these results prompted us to assess more precisely closeness of monogenic and GWAS IBD genes.

**Figure 1:**
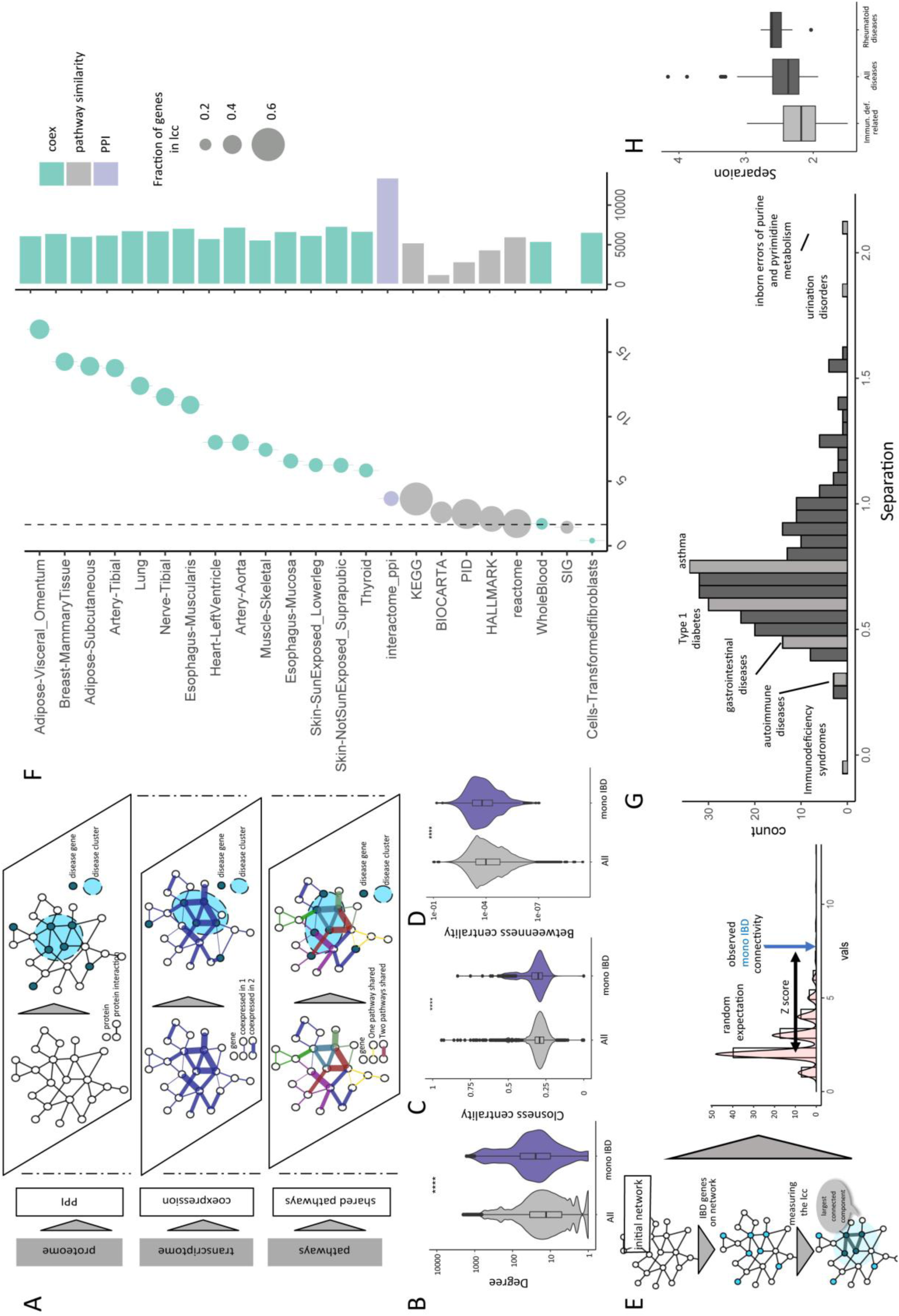
Monogenic inflammatory bowel disease on the interactome. a) Scheme of the multi-layered networks. The different layers represent protein-protein interaction, coexpression, and pathway similarity networks. b) Degree of monogenic IBD genes compared to all genes on the networks. c) Closeness centrality of monogenic IBD genes compared to all genes on the networks. d) Betweenness centrality of monogenic IBD genes compared to all genes on the networks. e) Explanation of the largest connected component (lcc). f) Monogenic IBD genes are significantly connected on the biological networks, measured by the z-score of the size of their lccs. g) Network-based separation of IBD genes from complex traits across the interactome. h) Network-based separation of monogenic IBD vs immunodeficiency and classical autoimmunity related terms. Significance of the lcc size (lcc z-scores) were calculated by comparing the lcc size of the monogenibc IBD genes to 1000 random permutations on the respective networks.

### Identification of putative novel monogenic IBD gene defects

A genetic pool of 638 unique SNPs associated with adult-onset (multifactorial) IBD was sourced from the GWAScatalog (https://www.ebi.ac.uk/gwas/) by quering for “inflammatory bowel disease”, “Crohn’s disease” and “ulcerative colitis” (Supplementary Figure 3A-B, Supplementary Table 2). Pathway enrichment revealed that the GWAS gene set was enriched for genes involved in TNF signaling, JAK-STAT signaling, T-cell receptor signaling, as well as prolactin cytokine signaling interleukin signaling and IFNγ signaling processes (Supplementary Figure 3C-D). As reported previously, some overlap was observed with GWAS genes connected with other diseases such as leprosy, and a broad spectrum of immune-mediated diseases, including psoriasis and ankylosing spondylitis^4^ (Supplementary Figure 3E), collectively highlighting the diverse molecular mechanisms and pathways likely involved in IBD pathogenesis. Interestingly, pathways in cancer was the top enriched term for IBD GWAS genes, reflecting the fact that the cancer-related gene group contains a considerable number of associations with major immune-related pathways but also likely pointing to the link between cancer and inflammation. Comparison of GWAS and monogenic IBD showed limited overlap at the gene level, perhaps because the majority of GWAS genes are functionally redundant (Supplementary Figure 3F). Interestingly, a high proportion of SNPs associated with Crohn’s disease and ulcerative colitis were shown to map to promoters or enhancers active in T cells^29^, suggesting an important contribution of adaptive immunity to the pathogenesis of polygenic forms of IBD as in monogenic IBD. To better assess whether and how polygenic and monogenic IBD might converge within functionally similar pathways, we compared them at the interactome level.

In total, 282 out of 573 GWAS genes and 53 out of 70 monogenic IBD genes were represented in the interactome, the rest being excluded for the downstream analyses. Collectively, they formed a highly significantly connected cluster of 101 genes (z=5.3, Supplementary Figure 4A-B). As expected, within the joint disease cluster, monogenic IBD was closer to inborn errors of immunity with IBD phenotype than to GWAS IBD (Supplementary Figure 4C). We hypothesized that the most biologically relevant GWAS genes within this cluster are those with similar, albeit less pronounced network characteristics as monogenic IBD genes in terms of connectivity and centrality. To systematically identify such genes, we designed a network-based pipeline, using the 70 monogenic IBD genes as starting point. We used a well-established diffusion method, specifically adapted to place less emphasis on highly connected network hubs and to rank genes with the more robust network closeness to monogenic IBD genes more highly^30,31^ (Figure 2A, Supplementary Figure 5A). Within the top 50 ranked GWAS genes (referred to as “top” genes from here forward), we found several genes associated with inborn errors of immunity (Figure 2B), illustrating the power of this approach in highlighting essential checkpoints of immune homeostasis. The top genes were further enriched in essential genes, as well as in genes associated with other orphan diseases (Figure 2B-C). Conversely, top genes were depleted in genes that may harbor loss of function mutations in the healthy population, so called “dispensable genes”, compared to GWAS genes farther away from the monogenic IBD genes (Figure 2D). While we only utilized interactome information to identify the top genes, they were also found to be central in the coexpression and pathway networks (Figure 2E-G). From the 43 top genes not yet implicated in a monogenic disease, we selected those with high disease-module cohesion, arriving at 14 GWAS genes associated with IBD (Figure 2H). Nodes with high disease-module cohesion are highly localized around the initial monogenic IBD gene set, therefore we hypothesized that these genes might be of particular functional importance in the pathogenesis of common forms of IBD and are also potential candidates for new monogenic disorders. To contextualize the putative functions of the top genes, we looked at the subnetwork defined by the top candidates and monogenic disease genes (Figure 3A). The subnetwork surrounding the top 14 candidates was enriched in proteins involved in global pathways such as pathways in cancer, PI3K-AKT signaling, T-cell receptor signaling, MAPK and RAS, Toll-like receptor signaling pathways, and the JAK/STAT pathway. Specific processes that were found to be strongly enriched included NK-cell mediated cytotoxicity and Fc gamma receptor mediated phagocytosis, highlighting the importance of genes involved in the regulation of inflammation, of the innate as well as of the adaptive immune system (Figure 3A, Supplementary Figure 5B-C).

**Figure 2:**
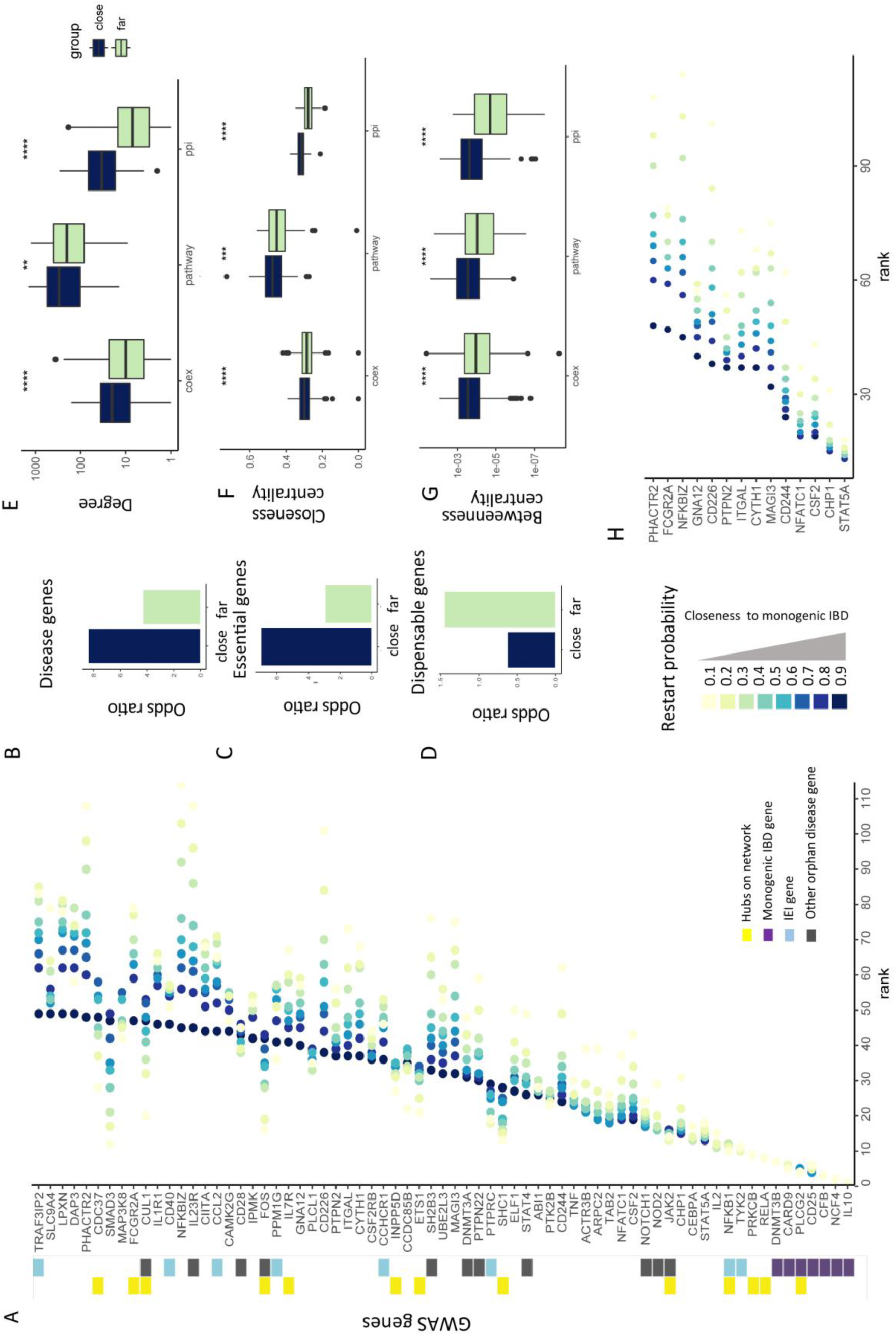
Identifying biologically relevant GWAS genes by disease-module cohesion. a) Ranking GWAS genes based on their closeness to monogenic IBD with a random walk with restart with changing restart probability. The best scoring genes receive the smallest ranks. Each colored dot shows a ranking of the respective gene in the random walk with a specific respart probability. All genes receiving a rank 50 or smaller are visualized. b-d) Enrichment of close GWAS genes as compared to far GWAS genes in essential, orphan disease and dispensable genes e-g) Degree, closeness centrality and betweenness centrality of close GWAS genes as compared to far GWAS genes on the different networks. h) The 14 top candidate genes with high disease-module cohesion. Significance of difference between networks based traits was measured by t-test (ns, non-significant, *p < .05, **p < .01, ***p < .001, ****p < .0001).

**Figure 3:**
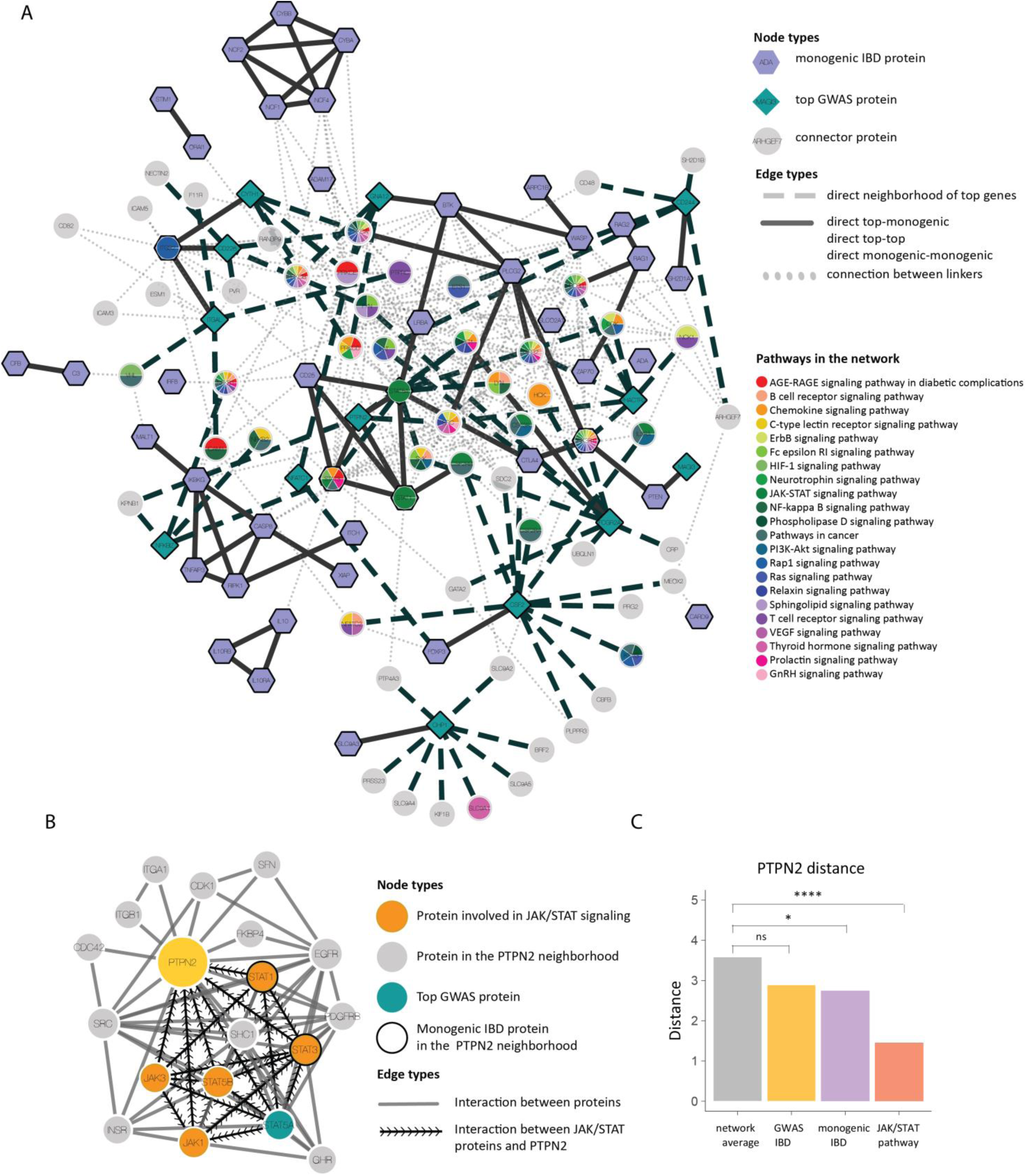
The network neighborhood of top genes. **a)** The close network neighborhood of the top candidates, monogenic IBD genes and close connectors on the interactome. Node shapes represent the different node types (top gene, monogenic IBD gene and connector) Nodes are colored according to pathway annotation. **b)** The subnetwork of PTPN2 on the interactome. **c**) Average shortest distance of PTPN2 from the monogenic IBD, GWAS IBD and the JAK/STAT pathway.

### Identification of a de novo PTPN2 mutation in a child with early onset intestinal autoimmunity

To validate our network-based approach, a cohort of genetically undiagnosed IBD patients (n=198) presenting with very early-onset IBD (VEO-IBD, defined by age of onset before 6 years), was screened for mutations in the the top 14 GWAS genes by whole-exome sequencing. This analysis identified a heterozygous *de novo*, nonsynonymous variant in exon 6 of the *PTPN2* gene (NM_002828.3 c.646T>G, chr4:765,966-775,965) (Supplementary Table 5) in a 5-year-old girl with severe autoimmune enteropathy since the age of three months. The child, born from heatlhy non-consanguineous Caucasian parents, presented with chronic secretory diarrhea, severe villous atrophy with prominent lymphocytic infiltrates in duodenal biopsies (Figure 4A), high titers of anti-AIE75 antibody but normal frequency of Foxp3^+^ CD4+ T cells (Supplementary Tables 6-7). Disease resisted over one year to treatment by azathioprine and tacrolimus but subsided when tacrolimus was replaced by sirolimus (Figures 4A-B). Further screening by targeted next-generation sequencing of 61 additional patients presenting also with autoimmune enteropathy did not reveal other pathogenic or rare heterozygous variants of *PTPN2*. Yet, a critical role of PTPN2 in immune regulation has been evidenced in *Ptpn2-/-* mice, which developed severe systemic and intestinal inflammation and autoimmunity immediately after birth, and died within 3 to 5 weeks^12,32–34^. Moreover, we observed that *PTPN2* is under significant purifying selection against loss-of-function (LoF) mutation with a pLI score of 1 and a missense ExAC z-score of 1.38, overall sustaining the hypothesis that PTPN2 plays a critical immunoregulatory role in humans and, therefore, that the observed *de novo* variant in PTPN2 may be a novel monogenic cause of intestinal inflammation in the affected child. To confirm this hypothesis, we analyzed the consequence of the observed missense mutation on the function of PTNP2.

**Figure 4:**
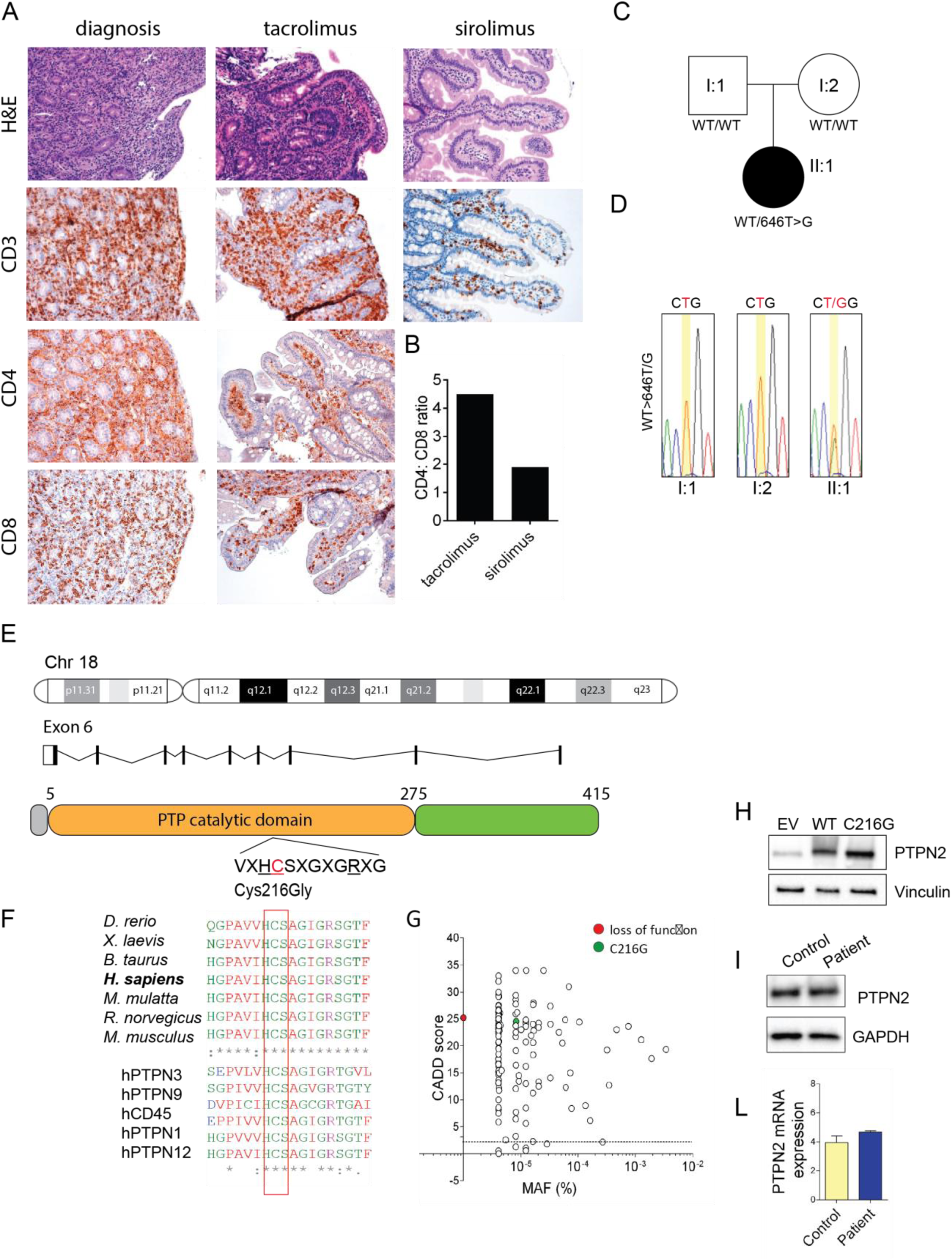
The *PTPN2* mutation occurs *de novo* and affects a highly conserved residue in the core of the catalytic site of the PTPN2. a) Hematoxylin and eosin staining of duodenal tissue obtained by biopsy showing villous blunting and CD3, CD4, CD8 staining indicating the presence of lymphocytic inflammatory infiltrates during the course of the disease. Original magnification, from left to right: H&E: 100x, 200x, 100x; CD3: 200x, 100x, 100x; CD4: 200x, 100x; CD8: 200x, 100x . b) CD4/CD8 ratio in peripheral blood before and after treatment. c) Pedigree showing a *de novo* heterozygous mutation in PTPN2, resulting in C216G amino acid substitution. The affected individual is denotated by a filled circle. WT, wild type. d) Sanger-sequencing electropherograms for the affected individual and parents. e) Chromosomal location, exon structure of the *PTPN2* gene and domain structure of the PTPN2 protein. f) Multiple alignments of PTPN2 orthologs from different species among species (top) and across the superfamily of tyrosine phosphatases (bottom) using the Clustal Omega software. The aminoacid affected by the mutation is boxed in red. Conserved residues are indicated as follow: full identity (*), similar characteristics (:) (>0.5 in the Gonnet PAM 250 matrix), weak similarities (.) (<0.5 in the Gonnet PAM 250 matrix). g) All nonsynonymous coding *PTPN2* variants previously reported in EXAC database and the PTPN2 missense variant identified here. The allele C216G is private to the family. The minor allele frequency (MAF) and combined annotation–dependent depletion (CADD) score of each variant are shown. The dotted line indicates the mutation significance cutoff (MSC) of PTPN2 with 95% confidence interval. h) PTPN2 expression in HEK293T cells overexpressing WT and C216G allele. i) PTPN2 protein and mRNA expression in patient and control-derived EBV cells.

### The C216G mutation impairs PTPN2 catalytic activity

*PTPN2* encodes a prototypic member of the PTP family of 38 enzymes that orchestrate dephosphorylation-dependent signal transduction (ref). The c.646T>G mutation detected in the exon 6 of *PTPN2* resulted in a p.Cys216Gly substitution (Figure 4C-E). This variant was absent in the Imagine Institute database of over 13,000 exomes and in public databases including the Exome Aggregation Consortium (EXAC) Database encompassing over 120.000 exomes or genomes (Figure 4G). *De novo* segregation pattern was confirmed by Sanger sequencing (Figure 4D). Strikingly, the affected amino acid, Cys216, is highly conserved across species as well as across other members of the superfamily of tyrosine-specific PTPs (Figure 4F). Accordingly, the p.Cys216Gly substitution was indicated to be damaging by all prediction tools (SIFT, Polyphen, Mutation Taster; Supplementary Table 5) and was associated with a high combined annotation-dependent depletion (CADD) score of 25.2 largely above the mutation significance cutoff of 3.13 for PTPN2 (Figure 4G), further supporting its damaging and deleterious effect^22,35^.

The c.646T>G mutation did not impact PTPN2 expression or stability. Thus, overexpression of the mutant allele by lentivirus-mediated gene transfer in HEK293T cells yielded a 45kDa PTPN2 protein in amounts comparable to the wild type (WT) protein (Figure 4H) and comparable level of endogenous PTPN2 mRNA and protein levels were found between patient’s and control derived lymphoblastoid cell lines (LCLs) (Figure 4I-L). In contrast, the mutation impaired PTPN2 enzymatic activity. Indeed, Cys216 is located in the core of the catalytic site and is part of the highly conserved ‘PTP signature motif’ (Figure 4E-F). This signature consists in the canonical sequence HC(X5)R and forms the phosphate-binding loop at the base of the active-site cleft in all members of the PTP family^36^. The cysteine residue initiates the first step of the catalysis by mounting a nucleophilic attack of the phosphate substrate via the thiolate group (Figure 5A). Replacement of cysteine with glycine should impair this catalytic step as glycine lacks the polar side chain to interact with the phosphorus atom of phospho-tyrosyl residue. To confirm the impact of PTPN2 C216G on catalytic activity, WT or mutant C-terminally tagged PTPN2 were transiently expressed in HEK293T cells, immunoprecipitated and de-phosphorylation of STAT1 or STAT3 peptides, that are putative substrates of PTPN2^37^ was measured. Compared to WT PTPN2, the mutant PTPN2 enzyme was totally deprived of catalytic activity (Figure 5B, Supplementary Figure 5A). As PTPN2 is known to inhibit Jak/STAT signaling^37^, we further tested whether the C216G mutation might result in hyperactivation of STAT3 signaling. Lentiviral constructs encoding WT or C216G-PTPN2 were used to transduce HEK293 stably expressing a luciferase reporter gene under the control of STAT3 transcriptional response elements (TRE). Following IL-6 stimulation, overexpression of WT PTPN2 led to significant downregulation of STAT3 TRE activity over control while the mutant form of PTPN2 failed to repress the reporter gene (Figure 5C).

**Figure 5:**
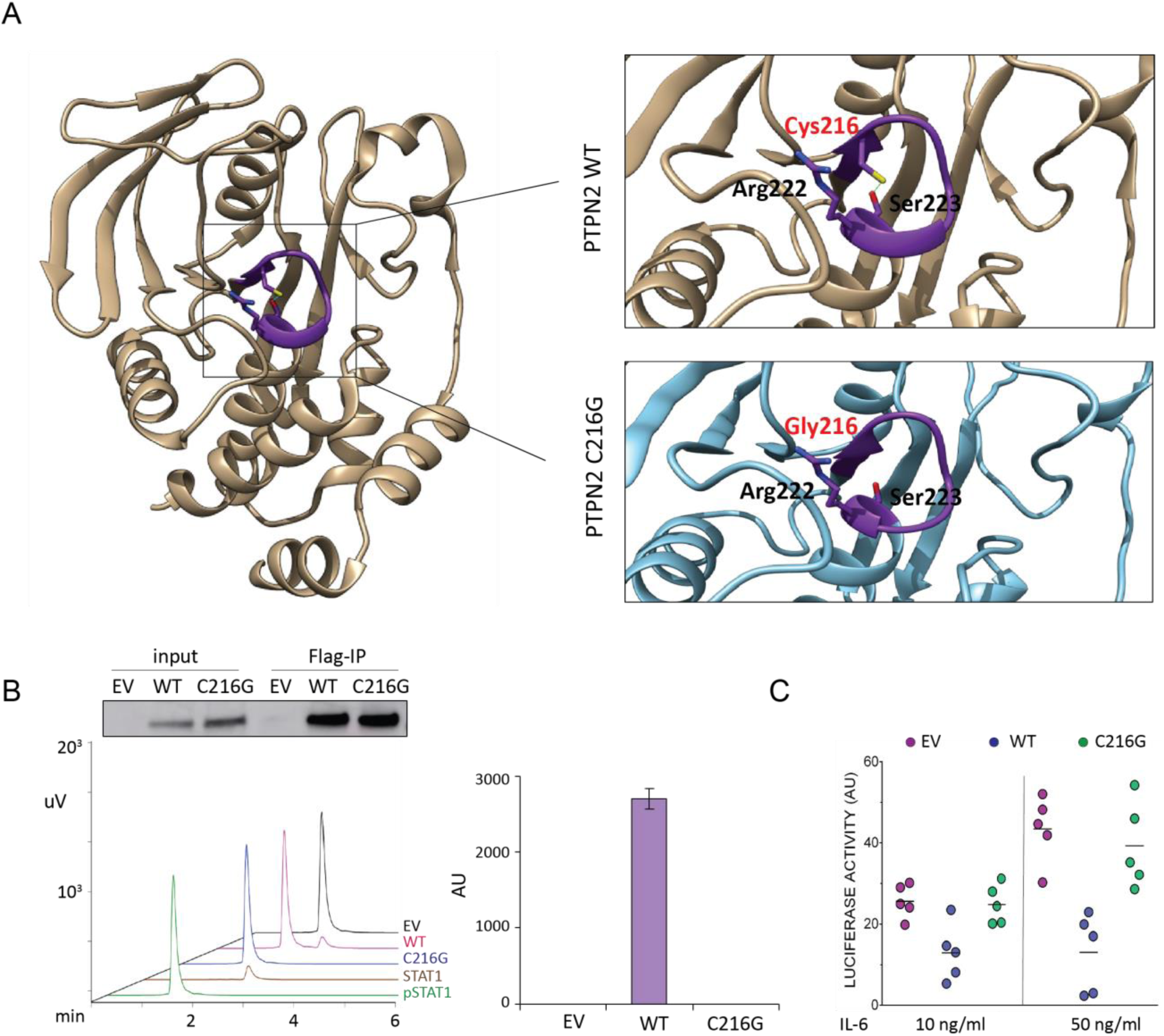
C216G is catalytically inactive. a) Mapping of Cys216 onto the crystal structure of human PTPN2 (Protein Data Bank (PDB) accession 1L8K). The position of the catalytic cysteine residue of PTPN2 (Cys216) and two other important residues also present in the PTP-Loop (Arg222 and Ser223) are shown in sticks. The zoomed-in region (with the structure rotated 90° toward the reader) shows the position of Cys216 (red), Arg222 and Ser223. The hydrogen bond between the lateral chains of Cys216 and Ser223 is shown with green dots. b) Determination of the tyrosine phosphatase activity of immunoprecipitated WT and C216G PTPN2 towards pStat1 was carried out by RP-UFLC analysis using a fluorescent tyrosine-phosphorylated Stat1 peptide. the RP-UFLC chromatograms are shown for control peptides (FAM-pStat1 and FAM-Stat1) and for the immunoprecipitates from empty vector, WT and C216G PTPN2 transfected HEK cells. PTPN2 activity toward pStat1 is shown in arbitrary units. Results were representative of experiments done in triplicate. c) STAT3 transcriptional activity following 48-hour activation with IL-6 (10 ng/mL) of HEK293T cells expressing STAT3-responsive luciferase and stably transduced with empty vector (EV), PTPN2 WT and C216G PTPN2. Results represent the mean of 5 independent experiments (ns, non-significant, *p < .05, **p < .01, one-way ANOVA).

### The PTPN2 C216G mutation results in haploinsufficiency

Whether PTPN2 works as a monomer or a homodimer is unclear^38^. It was therefore unsure whether expression of the mutated allele may or not impair the function of the residual normal protein. HEK293T cells were transiently cotransfected with plasmids expressing differentially tagged PTPN2, FLAG-PTPN2-FLAG and GFP-PTPN2. FLAG-immunoprecipitates contained large amounts of FLAG-PTPN2 but also some GFP-PTPN2, suggesting that, at least in overexpression setting, PTPN2 may homodimerize and, therefore, that the C216G allele could potentially act in a dominant negative manner (Figure 6A). Catalytic activity of a mix of recombinant WT and C216G PTPN2 at a 1:1 ratio was however comparable to the activity of recombinant WT PTPN2 alone (Figure 6B). Similarly, the capacity of transfected WT PTPN2 to repress the luciferase reporter in STAT3-TRE HEK293T cells stimulated by IL-6 was unmodified by cotransfection with C216G PTPN2 (Figure 6C). Collectively, these data indicate that the mutant allele does not exert a dominant negative effect and that the disease results rather from haploinsufficiency. This result is in keeping with PTPN2-pLI score of 1 that predicts that the loss of a single copy should be poorly tolerated and result in disease^39^.

**Figure 6:**
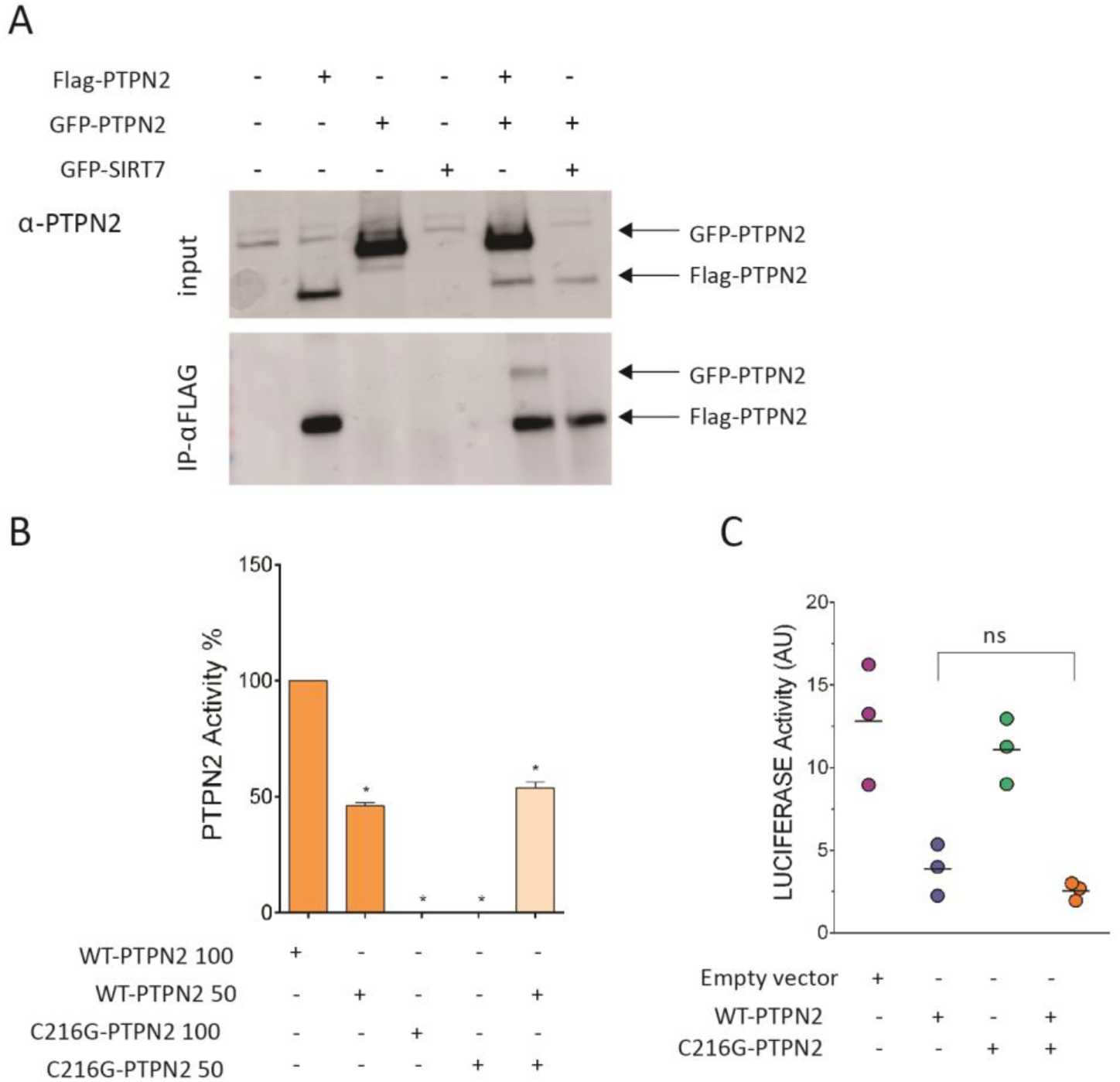
The C216G does not exert a dominant negative effect. a) co-immunoprecipitation of Flag- and GFP-tagged WT PTPN2 transfected into HEK293T cells at 1:1 vector ratio. b) PTPN2 activity assays were done with 200 ng and 100 ng of purified WT or C216G PTPN2 enzyme. Assays were also done with a mixture (1:1 ratio) of 100 ng WT PTPN2 and C216G PTPN2. Experiments were carried out in triplicate. c) STAT3 transcriptional activity following 48-hour activation with IL-6 (10 ng/mL) of HEK293T cells expressing STAT3-responsive luciferase and transfected with empty vector (EV), PTPN2 WT and C216G PTPN2 or both a 1:1 vector ratio. Results represent the mean of 3 independent experiments (ns, non-significant, *p < .05, **p < .01, one-way ANOVA).

### TCR signaling is preserved in human monoallelic loss-of-function *PTPN2* mutation

Studies in mice have shown that, *in vivo,* Ptpn2-deficient CD4 and CD8 T cells display reduced threshold of activation in the T-cell receptor (TCR) due to the lack of tyrosine de-phosphorylation of the Src family kinase family member Lck^40^. The impact of PTPN2 deficiency on TCR activation was assessed in T-cells derived from the patient and one healthy control upon TCR ligation by anti-CD3 antibody. In patient cells, global tyrosine phosphorylation of substrates of the TCR signaling cascade was not significantly different from that observed in control cells. Accordingly, we did not observe any change in Ca^2+^ mobilization (Supplementary Figure 7A), or in phosphorylation of ERK1/2 kinases downstream TCR activation between control and patient T blasts. Of note, in this setting, LCK was also equally phosphorylated (Supplementary Figure 7B). These results were recapitulated in PTPN2-depleted Jurkat cells (Supplementary Figure 7C), which, in contrast and as expected, displayed hyper-phosphorylation of STAT1 in response to IFN-γ (Supplementary Figure 7D). Overall these results suggest differences in the role of PTPN2 in human and mouse TCR activation, PTPN2 being likely redundant in this context in humans. Accordingly, tacrolimus, a calcineurin inhibitor which blocks Ca^2+^ dependent signaling downstream of TCR failed to cure the severe intestinal autoimmunity observed in the patient, further ruling out TCR hyperactivation as a driver of the disease.

### JAK/STAT signaling is hyper-activated in *PTPN2*-mutant patient cells

To investigate the functional consequences of PTPN2 haploinsufficiency in primary immune cells, we stimulated T lymphoblasts derived from the patient or from controls with IL-15 for 24 hours and measured STAT3 and STAT5 phosphorylation by immunoblotting. Patient’s T cells displayed increased STAT3 phosporylation compared to T cells from healthy donors (Figure 7A). Because limited blood sampling prevented extensive studies in T cells, this result was further confirmed in Epstein Barr virus (EBV)-transformed lymphoblastoid cell lines. In response to IL-21, patient’s EBV-B cells exhibited increased and more persistent phosphorylation of JAK1 and STAT3 compared to two controls (Figure 7B). Furthermore, phosphorylation of STAT5 and STAT1, which were not triggered by IL-21 in EBV-B cell lines from controls, was observed in EBV-B cells from the patient, pointing to alternative reprogramming of JAK-STAT signaling in patient cells.

**Figure 7:**
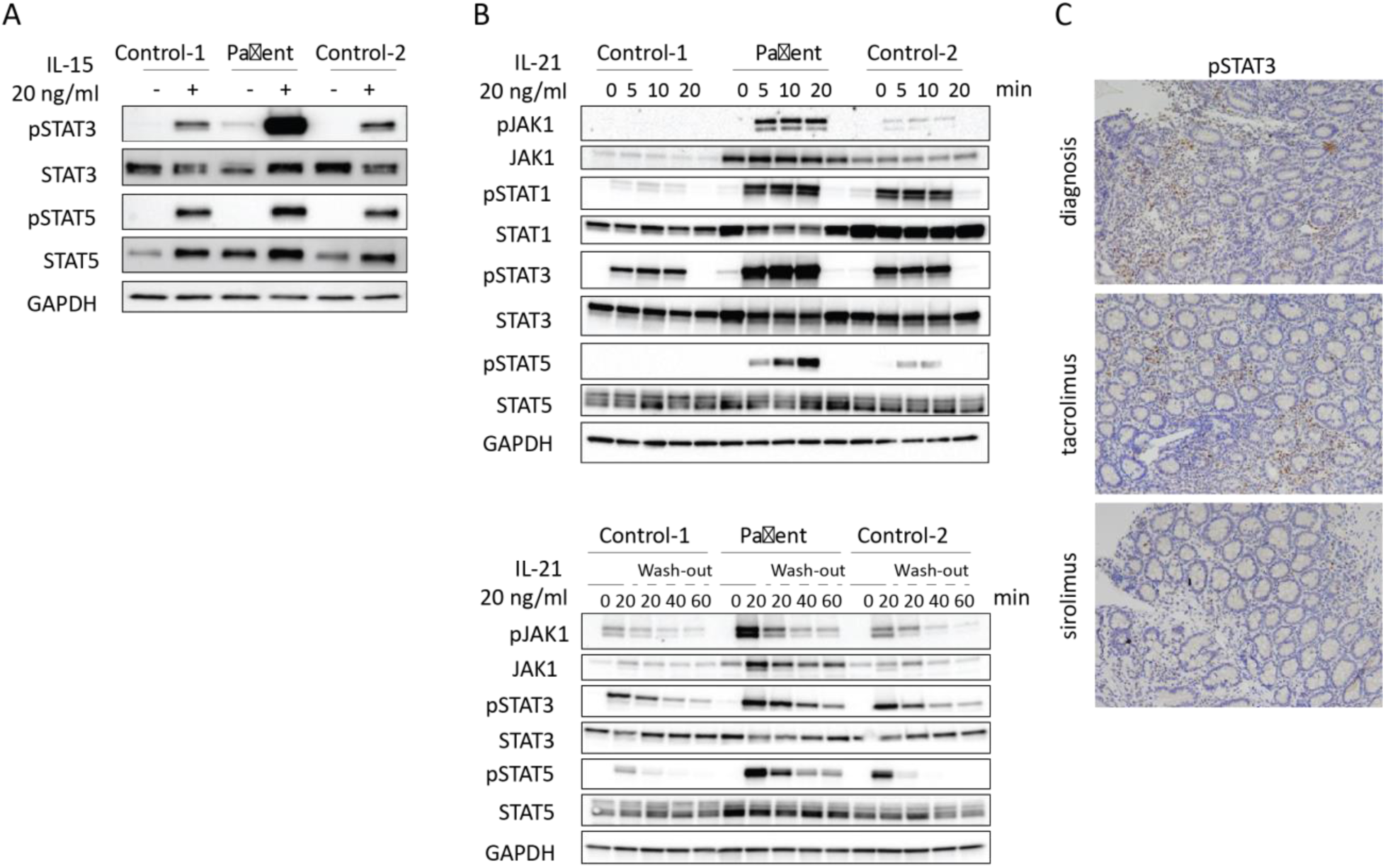
PTPN2 haploinsufficiency causes aberrant JAK/STAT signaling. a) Immunoblot of T blast cells obtained from two healthy controls and patient after 24 hours of stimulation in presence or absence of 20 ng/mL IL-15. Data representative of two independent experiments are shown. b) Immunoblot analysis of LCL obtained from two healthy controls and patient after 24 hours of stimulation in presence or absence of 20 ng/mL IL-21 for indicated time points (top panel) or after wash-out of the cytokine (bottom panel). Data representative of two independent experiments are shown. c) pSTAT3 staining of duodenal tissue during the course of the disease.

Finally, prominent STAT3 phosphorylation was detected in mononuclear cells infiltrating intestinal biopsies of the C216G patient at time of disease activity (Figure 7C). In contrast, STAT3 phosphorylation was not detected in epithelial cells, a result in line with the lack of intestinal inflammation in mice with conditional *Ptpn2* deletion in enterocytes ^14^. Overall these findings, are reminiscent of observations made in the inflamed colons of mice carrying CD4-specific deletion of *Ptpn2*^13^. They support the hypothesis that the mutation licences excessive *in vivo* activation of the JAK1-STAT3 pathway in lymphocytes and drives intestinal pathology (Figure 7C).

## DISCUSSION

IBD generally arise from a complex interaction of genetic and environmental factors. Although animal models, eQTL, and epigenetic fine mapping^29,41^ have revealed valuable insights into the role of certain SNPs and GWAS genes, it remains challenging to establish causal links with the pathobiology of IBD. IBD can also manifest as monogenic, early-onset diseases, thereby providing an unique opportunity to study the roles of susceptibility genes in clear genotype-to-phenotype pathobiology. In contrast to GWAS which mainly underscore the contribution of innate immune pathways, genes underlying monogenic IBD tend to affect molecular mechanism and cell types involved in adaptive immunity-related processes (Supplementary Figure 1C). This duality behind the pathomechanism of IBD remains unexplored today. We collected multiple biological networks that reflect diverse mechanisms known to be involved in IBD pathobiology, such as physical protein-protein interactions or transcriptional regulation. We find that monogenic IBD genes form significant disease clusters and are characterized by central positions within networks ranging from the transcriptome to proteome and functional levels (Figure 1B-D, Supplementary Figure 2A). The importance of network centrality as an indicator of gene essentiality has been shown for multiple species^42^. Along the same lines, according to the recently proposed “omnigenics” model of diseases^43,44^, mutations within central core genes will impact regulatory networks more strongly than variations within less central genes, thus inducing a disease phenotype more directly. Therefore, to explore the hematopoietic component of IBD and identify essential links connecting the innate and adaptive processes underlying bowel inflammation, we used genes that underlie monogenic IBD and PID with IBD as starting point of our analysis. To identify GWAS genes potentially playing such central roles for IBD and the IBD disease-module on networks, we developed a network-based pipeline custom-tailored for IBD. Building on previous approaches for utilizing known functional relationships among genes to contextualize and rank candidate genes according to their proximity to known disease-associated genes^30,45,46^, we incorporated IBD specific molecular interaction types, and introduce disease-module cohesion as the means to select our top candidates. The network neighborhood of the final top candidates of our pipeline was enriched with IBD-associated pathways, including the JAK/STAT pathway, as well as with cancer-related pathways, highlighting the emerging parallel between chronic inflammation and malignancy^47^.

We have identified 14 GWAS genes which were not yet associated with monogenic diseases but had high disease-module cohesion. By screening a cohort of VEO-IBD patients, we identified a *de novo* mutation resulting in *PTPN2* haplo-insufficiency in a child with intestinal autoimmunity of very early onset. The missense mutation replaced the highly conserved cysteine of the ‘PTP signature motif’ that is indispensable for the catalytic activity of Class I-III phosphatases^36,48^ by a glycine residue. Accordingly, the C216G PTPN2 mutant allele was totally deprived of phosphatase activity. In keeping with evidence that PTPN2 negatively regulates the activation of the JAK-STAT pathway by dephosphorylation of phospho-tyrosine residues of JAK and STAT molecules^37^, we showed that *PTPN2* haploinsufficiency impaired the regulation of the JAK-STAT pathway in lymphocytes. In contrast with data in mice with complete inactivation of Ptpn2^40^ but in keeping with the lack of therapeutic efficacy of calcineurin inhibitors, T-cell receptor signaling was not detectably enhanced. GWAS have repetitively associated intronic single nucleotide polymorphisms (SNP)s in the *PTPN2* locus with chronic inflammatory and autoimmune diseases^49,50^. Whether or not these common variants impact PTPN2 expression and function remain unclear^51^. Yet, our biological network analysis allowed to extract the JAK/STAT as a disease relevant pathway out of the GWAS data and the identification of human *PTPN2* haploinsufficiency as a monogenic cause of autoimmunity further validated the relevance of this pathway in intestinal immunoregulation. We have recently shown that Ruxolitinib (JAK1/2 inhibitor) represents a therapeutic option for severe enterocolitis associated with *STAT3* GOF mutations^52^. Our present data indicate that JAK inhibitors may be an alternative to the treatment by rapamycin that succeeded in controling intestinal inflammation in our patient. More generally they provide genetic-based rationale for using drugs targeting the JAK/STAT pathway in IBD, some of which have been recently approved. Of note, deletion of *PTPN2* gene has been identified in T-cell acute lymphoblastic leukemia^53,54^. Moreover, our recent work suggests that GOF somatic mutations in *JAK1* and *STAT3* are main drivers of intestinal malignant lymphoproliferation complicating celiac disease, an autoimmune-like enteropathy driven by dietary gluten. Our observation of an autoimmune intestinal disease driven by a germinal LOF mutation in *PTPN2* provides a novel striking example of the very close link between autoimmunity and malignant lymphoid transformation and emphasizes the power of human genetics for mapping crucial pathways and defining their level of redundancy in intestinal homeostasis.

## Acknowledgments

We thank our patients and their families. We thank Dr Bosi for anti AIE75 serum measurement.

## Supplementary Materials

### Supplementary Methods

Supplementary Figure 1. Genetics of pediatric inflammatory bowel disease.

Supplementary Figure 2. Network-based properties of monogenic IBD.

Supplementary Figure 3. Genetics of polygenic/adult inflammatory bowel disease.

Supplementary Figure 4. GWAS and monogenic IBD on the interactome.

Supplementary Figure 5. GWAS prioritization and top gene enrichment.

Supplementary Figure 6. PTPN2 activity toward pSTAT3.

Supplementary Figure 7. TCR signaling is preserved in PTPN2 deficient condition.

Supplementary Table 1. Monogenic IBD genes.

Supplementary Table 2. GWAS SNPs and GWAS IBD genes (See: Supplementary Document 1).

Supplementary Table 3. All ranked GWAS genes (See: Supplementary Document 2).

Supplementary Table 4. Top ranked GWAS genes (See: Supplementary Document 3).

Supplementary Table 5. Immunoglobulins and serum antibody response of PTPN2-deficient patient.

Supplementary Table 6. Autosomal recessive (AR) and *de novo* variants identified by WES in PTPN2-deficient patient.

Supplementary Table 7. Lymphocyte subsets of P1.

