## supplementary material for "Network Analysis of Inflammatory Bowel Disease Reveals PTPN2 As New Monogenic Cause of Intestinal Inflammation"

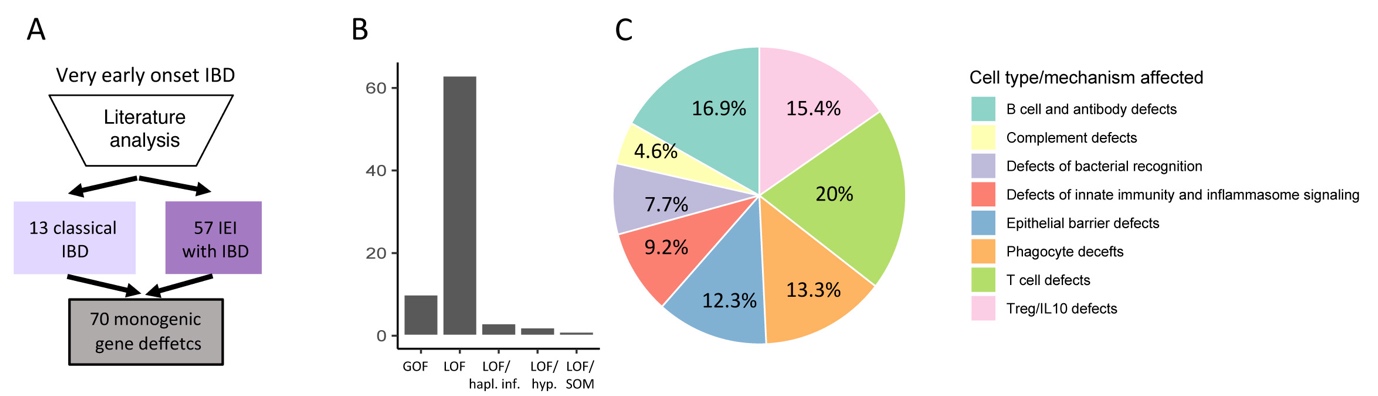

**Supplementary Figure 1: Genetics of pediatric inflammatory bowel disease.** a) 70 monogenic gene defects can underlie bowel inflammation. b) Distribution of functional consequences of gene defects underlying monogenic IBD (GOF: gain-of-function, LOF: loss-of-funtion, LOF/hapl: loss of function and haploinsufficiency, LOF/hyp loss-of-function hypomorphic, LOF/som: loss-of-function somatic). c) Types of monogenic gene defects underlying IBD according to affected process or cell type.

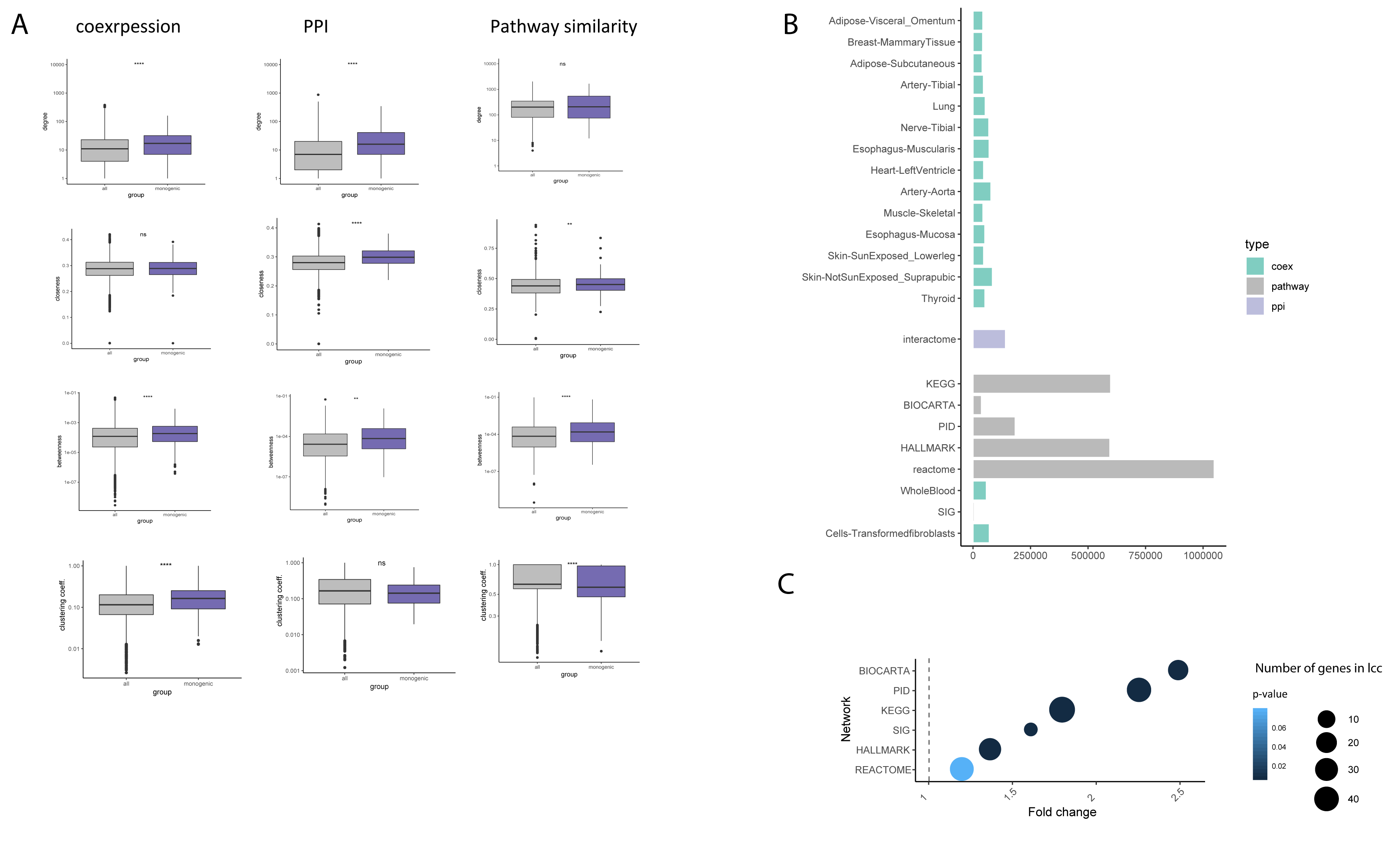

**Supplementary Figure 2: Network-based properties of monogenic IBD.** a) Number of edges per network. b) Enrichment in edge weights within the lcc of monogenic IBD genes on pathway similarity networks. c) degree, closeness centrality, betweenness centrality and clustering coefficiency of monogenic IBD genes compared to all other genes on the networks. Significance of the differences was calculated by t-test (ns, non-significant, *p < .05, **p < .01, ***p < .001, ****p < .0001) .

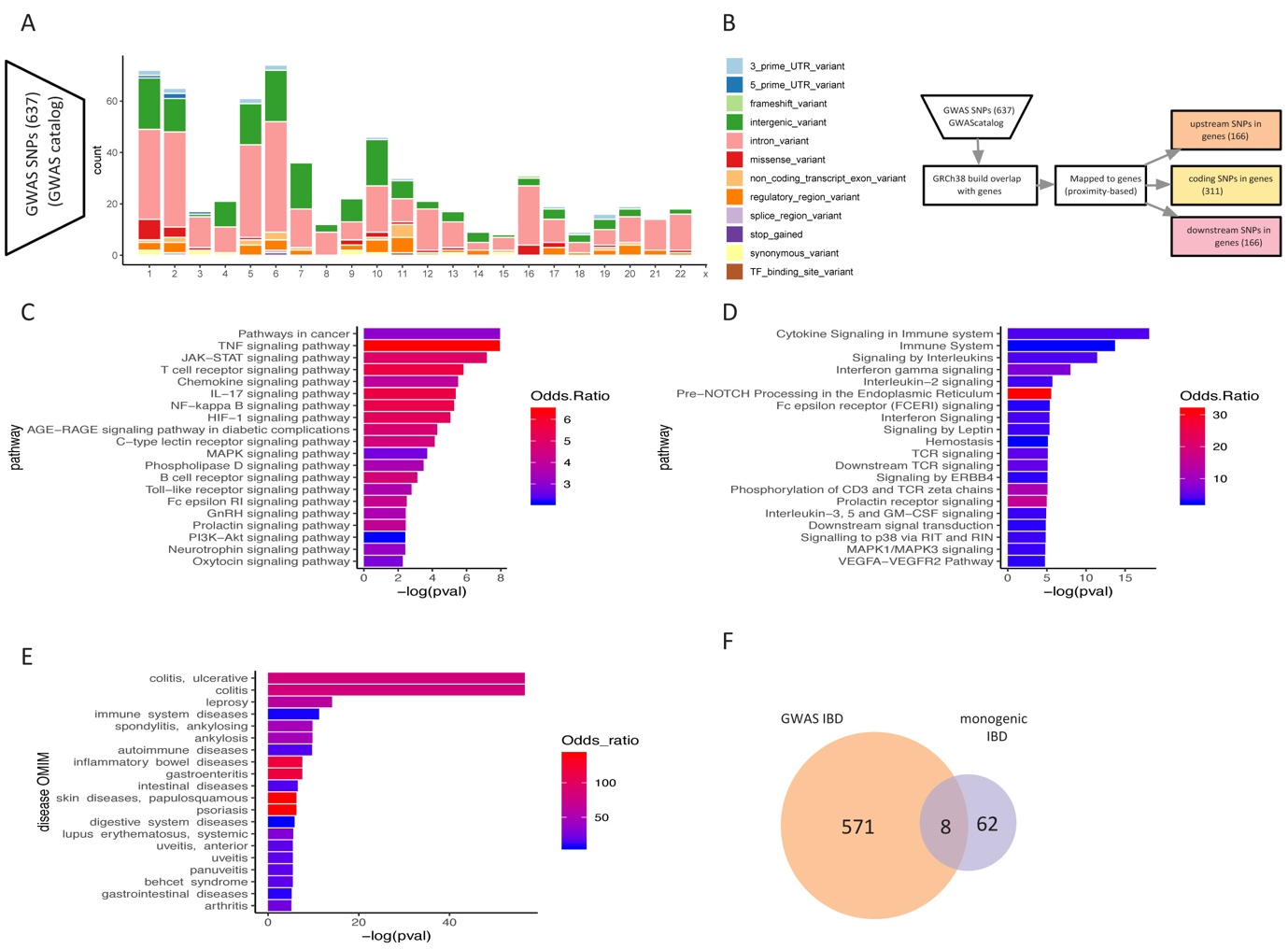

**Supplementary Figure 3: Genetics of polygenic/adult inflammatory bowel disease.** a) Distribution of significant SNPs associated with IBD across the genome and across functional context b) mapping of IBD associated GWAS genes c) global KEGG pathway enrichment of GWAS genes D) Enrichment of Reactome pathway terms in GWAS genes e) Enrichment of GWAS genes in OMIM disease terms f) overlap of monogenic and GWAS genes. F-H) mean intolerance of loss-of-function, missense and synonymous variation.

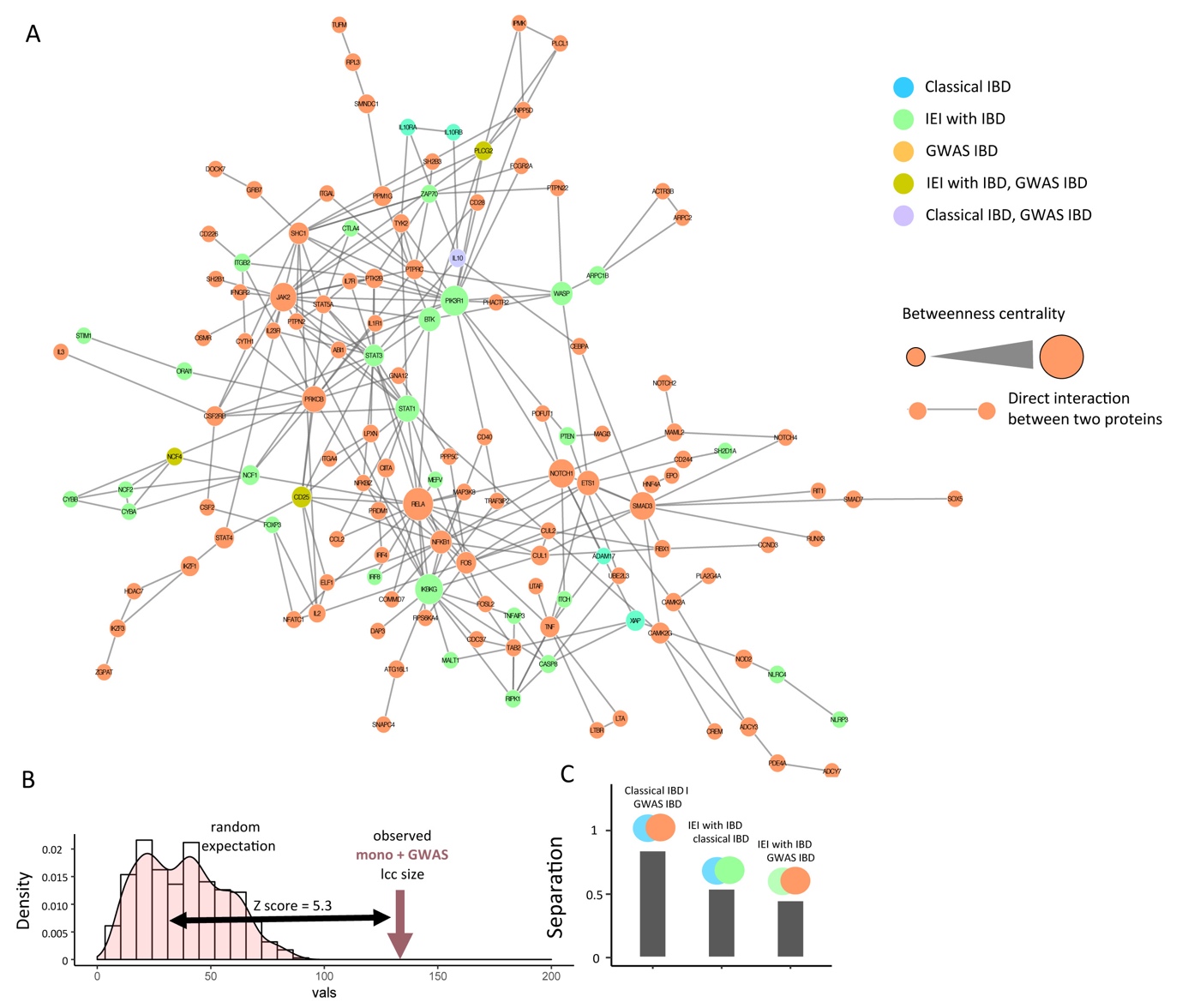

**Supplementary Figure 4: GWAS and monogenic IBD on the interactome** a) The disease module defined by GWAS and monogenic IBD gene on the interactome. b) The size and significance of lcc defined by the 70 monogenic IBD genes and 101 GWAS IBD genes is significantly connected on the interactome. c) Network-based separation of classical monogenic IBD, IEI with IBD and GWAS IBD, within the joint disease cluster on the interactome*.* IEI: inborn errors of immunity.

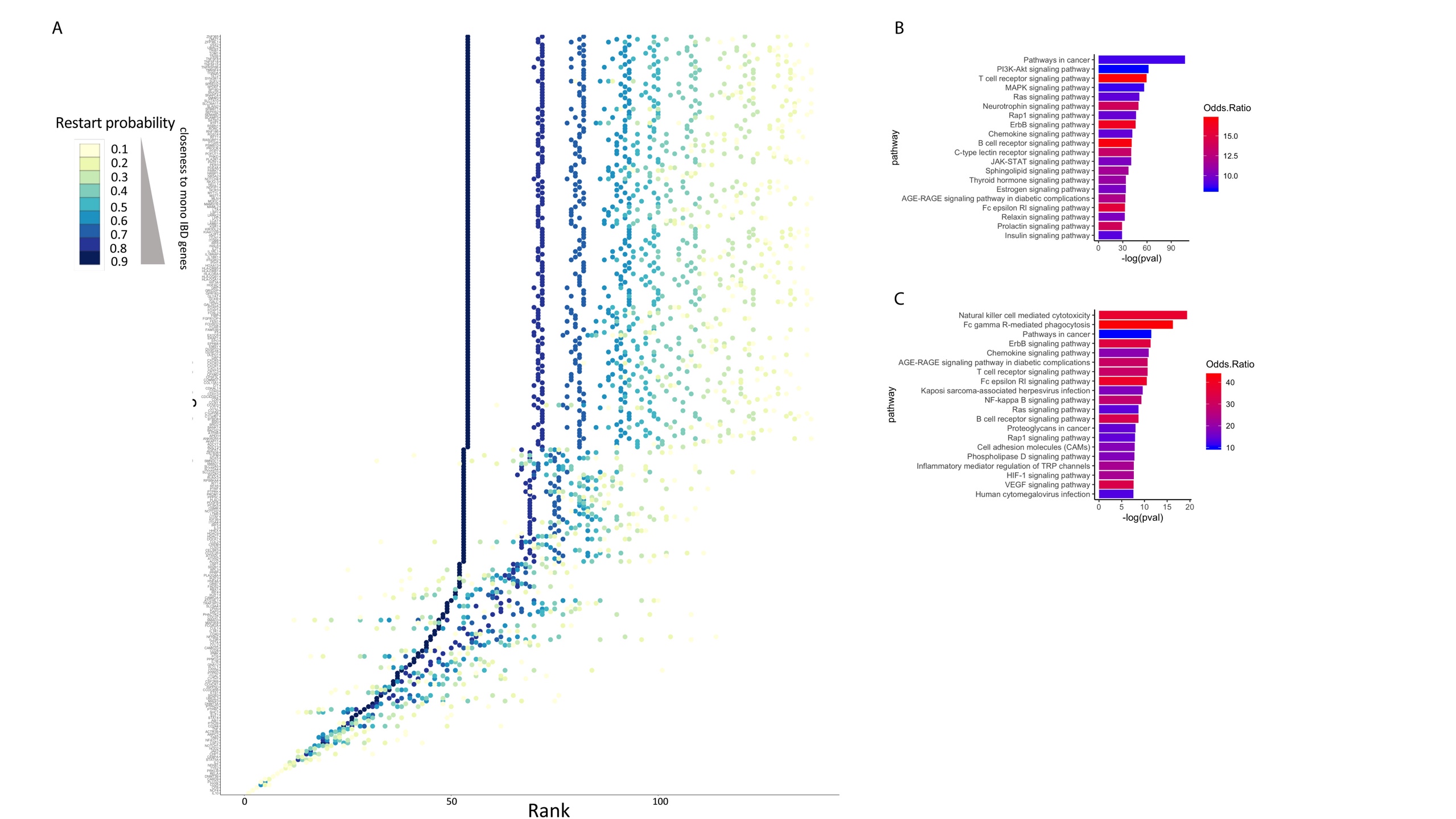

**Supplementary Figure 5: GWAS prioritization and top gene enrichment** a) All IBD GWAS genes on the interactome ranked by the iterative random walk approach. b) global KEGG pathway enrichment of the interactome neighborhood of the top genes (n=681 nodes). c) KEGG enrichment of the close neighborhood (n=60 nodes as displayed in Fig. 3A). Fisher's exact test was used to obtain p-value and odds ratio.

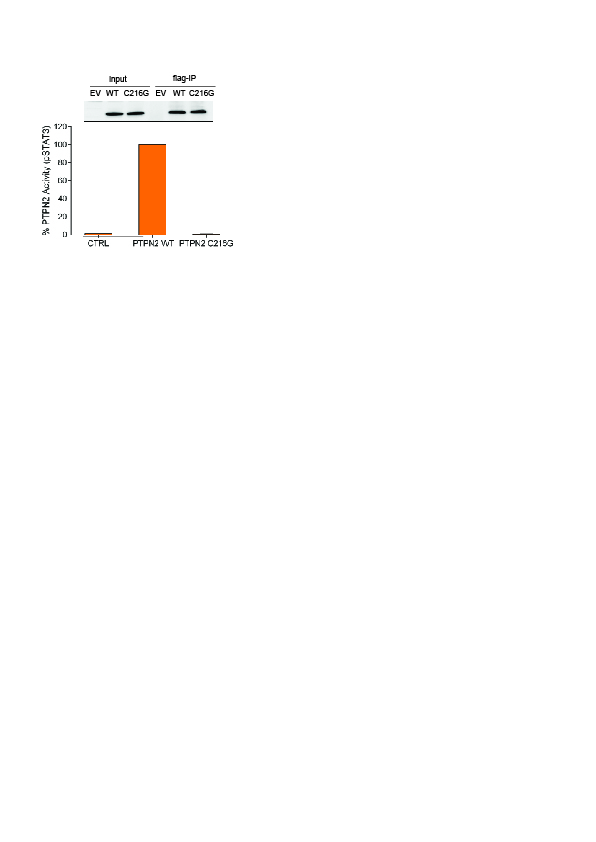

**Supplementary Figure 6: PTPN2 activity in immunoprecipitates using FAM-pStat3 and RP-UFLC.** Tyrosine phosphatase activity of immunoprecipitated WT and C216G PTPN2 towards pStat3 is carried out by RP-UFLC analysis using a fluorescent tyrosine-phosphorylated Stat1 peptide as described previously (Duval et al., 2015). PTPN2 activity towards pStat3 peptide is shown as percentage of the activity of WT enzyme.

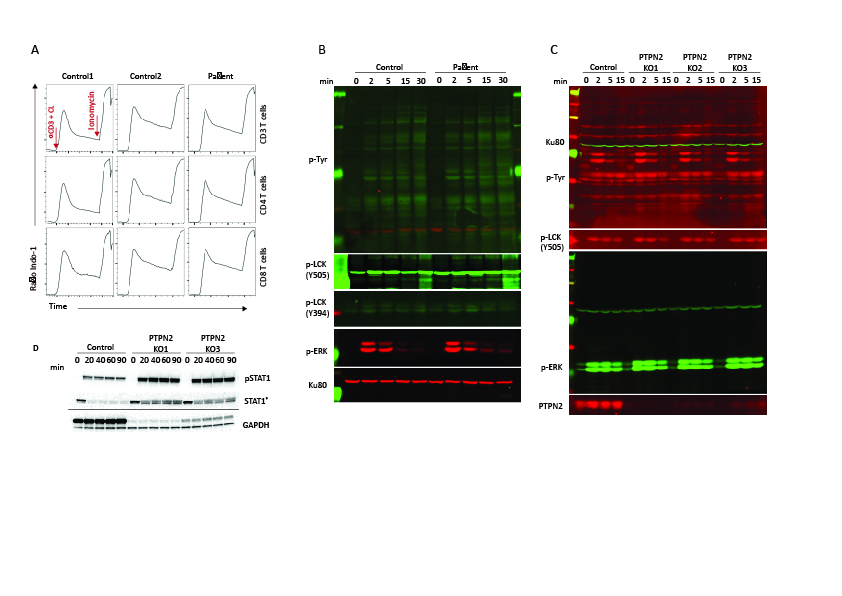

**Supplementary Figure 7: Calcium flux, global tyrosine phosphorylation and ERK1/2 phosphorylation is unaffected in activated PTPN2‐deficient T cells.** a**)** Flow cytometry analyses of Ca^2+^ flux in T‐cell blasts of a control donor and patient loaded with the Ca2+‐sensitive fluorescent dye Indo‐1, stimulated with anti‐CD3 antibody (first arrow) crosslinked with rabbit anti‐mouse antibody (CL) and then incubated with ionomycin. **b-c)** Immunoblot showing the global tyrosine phosphorylation, pERK, pLCK in T‐cell blasts from a control donor and Patient or in PTPN2-depleted Jurkat cells stimulated with anti‐CD3 antibodies at the time points indicated. **d)** Enhanced responsiveness to IFN-g stimulation as displayed by increased phosphorylation status of STAT1 in PTPN2-depleted Jurkat cells compared to control cells. *anti-STAT1 antibody preferentially recognize the non-phosphorylated form of Stat1.

**Table S1: Monogenic IBD genes**

| gene | year_of_first_publication | kind |
| --- | --- | --- |
| XIAP | 2006 | mono_IBD |
| IL10, IL10RA,IL10RB | 2009 | mono_IBD |
| TTC37 | 2010 | mono_IBD |
| ADAM17 | 2011 | mono_IBD |
| TTC7A | 2013 | mono_IBD |
| IL21 | 2014 | mono_IBD |
| DUOX2,NOX1 | 2015 | mono_IBD |
| TRIM22 | 2016 | mono_IBD |
| ANKZF1,CD55 | 2017 | mono_IBD |
| ALPI | 2018 | mono_IBD |
| C3 | 1972 | mono_IBD |
| NLRP3 | 1974 | mono_IBD |
| ADA | 1975 | mono_IBD |
| ITGB2 | 1981 | mono_IBD |
| CYBB | 1989 | mono_IBD |
| CYBA, IL2RG, PEPD | 1990 | mono_IBD |
| NCF1 | 1991 | mono_IBD |
| BTK | 1993 | mono_IBD |
| ZAP70 | 1994 | mono_IBD |
| NCF2 | 1995 | mono_IBD |
| ORAI1, RAG1, RAG2 | 1996 | mono_IBD |
| IL2RA;MEFV,PTEN | 1997 | mono_IBD |
| SH2D1A,DKC1 | 1998 | mono_IBD |
| DNMT3B, MVK | 1999 | mono_IBD |
| AICDA | 2000 | mono_IBD |
| DCLRE1C, FOXP3,IKBKG,LIG4,STAT1 | 2001 | mono_IBD |
| WAS | 2003 | mono_IBD |
| STAT3 | 2007 | mono_IBD |
| G6PC3,NCF4,STIM1,STXBP2 | 2009 | mono_IBD |
| ITCH,KRAS | 2010 | mono_IBD |
| ZBTB24 | 2011 | mono_IBD |
| LRBA,PLCG2 | 2012 | mono_IBD |
| CARD9,CFB,RTEL1 | 2013 | mono_IBD |
| CTLA4,NLRC4,TRNT1 | 2014 | mono_IBD |
| DOCK2,MASP2,NFAT5,PIK3R1,POLA1 | 2015 | mono_IBD |
| SLC9A3,TNFAIP3 | 2016 | mono_IBD |
| ARPC1B,IRF8,MALT1,SLCO2A1 | 2017 | mono_IBD |
| RIPK1, CASP8 | 2018 | mono_IBD |

**Table S2**. GWAS SNPs and GWAS IBD genes (See: Supplementary Document 1).

**Table S3.** All ranked GWAS genes (See: Supplementary Document 2).

**Table S4.** Top ranked GWAS genes (See: Supplementary Document 3).

**Table S5: Immunoglobulins and serum antibody response of P1**

|  | **7 mo of age** | | **1 yr 5 mo of age** | | **2 yr of age** | | **2 yr 7 mo of age** | |
| --- | --- | --- | --- | --- | --- | --- | --- | --- |
|  | **Patient** | **Reference range** | **Patient** | **Reference range** | **Patient** | **Reference range** | **Patient** | **Reference range** |
| IgG (g/Liter) |  |  |  |  | 6.15 | 4.24-10.51 |  |  |
| IgA (g/Liter) |  |  |  |  | 1.05 | 0.14-1.23 |  |  |
| IgM (g/Liter) |  |  |  |  | 1.49 | 0.18-1.68 |  |  |
| IgE (kIU/Liter) |  |  |  |  | < 2 | < 40.3 |  |  |
| anti-AIE75 (dpm) | **575** | <150 | **467** | <150 |  |  | **203** | <150 |
| anti-TPO (UI/ml) |  |  |  |  | **9.7** | < 9.0 | **9.1** | <9.0 |
| anti-TG (UI/ml) |  |  |  |  | < 2.0 | < 4.0 | <2 | <4.0 |
| Anti-GAD65 (dpm) |  |  |  |  | 33 | < 90 |  |  |
| Anti-IA2 (dpm) |  |  |  |  | 2 | < 55 |  |  |
| Anti-ZNT8 (dpm) |  |  |  |  | 6 | < 150 |  |  |
| Anti-DNA (UI/ml) |  |  |  |  |  |  | 3.5 | <7 |
| Anti-ANA |  |  | **1/200** |  |  |  | **1/100** |  |

**Table S6: autosomal recessive (AR) and *de novo* variants identified by WES in PTPN2-deficient patient.**

| **AR** | **Gene** | **Variant** | **Consequence** | **Sift** | **Mut taster** | **Polyphen** | **CADD** | **DB freq** |
| --- | --- | --- | --- | --- | --- | --- | --- | --- |
|  | ***SRRM2*** | rs144257955 | Thr2513Ala | Tolerated | Polymorphism | Benign | 10 | 0.299% |
|  | ***LY75*** | rs116058499 | Arg763Gln | Tolerated | Disease causing | Probably damaging | 27 | 0.217 % |
|  |  | rs113023766 | Met1308Val | Tolerated | Polymorphism | Benign | 23.9 | 0.137% |
|  | ***TDRD6*** | rs139660386 | Lys1432Arg | Tolerated | Polymorphism | Benign | 0 | 0.236% |
|  |  | rs144889394 | Ala491Val | Deleterious | Disease causing | Probably damaging | 24.3 | 0.020% |
|  | ***COL22A1*** | rs766709365 | Asp156Glu | Deleterious | Disease causing | Probably damaging | 14.7 | 0.005 % |
|  |  | rs770249679 | Arg407His | Deleterious | Disease causing | Probably damaging | 23.7 | 0.001 % |
|  | ***FOXD4L6*** | 9_69200873_G_A | Pro247L | Tolerated | Not scored | Benign | 8.3 | 0.265 % |
|  |  | rs2989709 | Pro416Arg | Tolerated | Not scored | Benign | 0 | 0.210 % |
|  | ***DOCK6*** | rs778570965 | Asp1804Ala | Deleterious | Disease causing | Benign | 25.1 | 0.001 % |
|  |  | rs183060698 | Val45Ile | Tolerated | Disease causing | Benign | 19.8 | 0.180 % |
| ***de novo*** | **Gene** | **Variant** | **Consequence** | **Sift** | **Mut taster** | **Polyphen** | **CADD** |  |
|  | ***NBPF1^#^*** | rs753338419 | Asn497Ile | Deleterious | not scored | Probably damaging | 22.7 | 0.011 % |
|  | ***NES^#^*** | 1_156640666_A_C | Val1105Gly | Deleterious | not scored | Benign | 11.5 | 0.065 % |
|  | ***HRTC1^#^*** | 9_35906602_C_CCACCA | frameshift | not scored | not scored | not scored | - | // |
|  | ***DNASE1L^#^*** | 16_2287508_G_C | Arg150Pro | Tolerated | not scored | Benign | 0.1 | 0.099 % |
|  | ***CSH1*** | rs1130686 | Pro3Thr | Tolerated | Polymorphism | not scored | 0 | // |
|  | ***PTPN2*** | 18_12817214_A_C | Cys216Gly | Deleterious | disease causing | Probably damaging | 25.2 | // |
|  | ***FUT3^#^*** | 19_5844663_A_G | Leu63Pro | Tolerated | not scored | BENIGN | 15.1 | 0.099 % |
|  | ***ALPPL2^#^*** | rs139018608 | Pro22Gln | Deleterious | Disease causing | Probably damaging | 22.5 | 0.779 % |
|  | ***USP36^#^*** | rs143011665 | Arg1006Cys | Tolerated | Polymorphism | Benign | 12 | 0.270 % |
|  | ***SKA3^#^*** | 13_21729290_T_TCAGTTTTCTTTGTTGCTGACATCTCGGATGTTCTGTCCATGTTTAAGGAACCTTTTA | ins/ splice acceptor donor | Not scored | Not scored | Not scored | - | 0.043 % |

^#^ the variant was considered unlikely causative as in our inhouse database was found in at least 15 individuals

**Supplementary Table S6:** Lymphocyte subsets in P1.

|  | 6 mo of age | | 2 yr of age | |
| --- | --- | --- | --- | --- |
|  | **Patient** | **Reference range** | **Patient** | **Reference range** |
| **CD3^+^ T cells (%)** | 71.8 | 48.0- 72.4 | 78 | 56- 75 |
| **CD3^+^ CD4^+^(%)** | 56 | 25.6- 52.5 | 45 | 28- 47 |
| **HLA-DR (%)** | 5.9 | 1.4- 17.6 |  |  |
| **CD45RA^+^ CCR7^+^** | 85.4 | 54.4– 88.6 |  |  |
| **CD45RA^+^ CCR7^+^ CD31^+^** | 66.4 | 49.9– 79 |  |  |
| **CD45RA^-^ CCR7^+^ (%)** | 7.5 | 6.4– 25.5 |  |  |
| **CD45RA^-^ CCR7^-^(%)** | 2.8 | 3.1– 16.8 |  |  |
| **CD45RA^+^ CCR7^-^(%)** | 4.4 | 0.7– 5.9 |  |  |
| **FOXP3^+^ CD217^lo^ CD25^+^(%)** |  |  | 4.4 | 4.5- 7.5 |
| **CD3^+^ CD8^+^ (%)** | 12.3 | 12.4– 20.3 | 23 | 16- 30 |
| **HLA-DR (%)** | 6.3 | 2.1– 52.0 |  |  |
| **CD45RA^+^ CCR7^+^(%)** | 82.8 | 36.6– 89.2 |  |  |
| **CD45RA^-^ CCR7^+^(%)** | 1.9 | 2.2– 8.3 |  |  |
| **CD45RA^-^ CCR7^-^(%)** | 3.4 | 5.4– 38.8 |  |  |
| **CD45RA^+^ CCR7^-^(%)** | 12 | 3.8– 16.2 |  |  |
| **CD 56+ CD16+ NK cells (%)** | 9.6 | 3.5- 23.5 | 8 | 4- 17 |
| **CD19 B cells (%)** | 17.2 | 10.7- 43.9 | 14 | 7.6- 28.2 |
| **CD38 ^dim/lo^ CD21^hi^ CD10^-^ CD27^-^ naive (%)** | 58.1 | 41.1- 65.1 |  |  |
| **IgD^-^ CD27^+^ class switched memory (%)** | 1 | 0.7- 5 | 1.6 | 2.7- 12.5 |
| **IgD^+^ CD27^+^ non switched memory (%)** | 3.7 | 3.6 -7 | 9 | 4.6- 16.3 |
